# Behavioral evolution drives hindbrain diversification among Lake Malawi cichlid fish

**DOI:** 10.1101/467282

**Authors:** Ryan A. York, Allie Byrne, Kawther Abdhilleh, Chinar Patil, J. Todd Streelman, Thomas E. Finger, Russell D. Fernald

## Abstract

The evolutionary diversification of animal behavior is often associated with changes in the structure and function of nervous systems. Such evolutionary changes arise either through alterations of individual neural components (“mosaically”) or through scaling of the whole brain (“conceitedly”). Here we show that the evolution of a specific courtship behavior in Malawi cichlid fish, the construction of mating nests known as bowers, is associated with rapid, extensive, and specific diversification of orosensory, gustatory centers in the hindbrain. We find that hindbrain volume varies significantly between species that build pit (depression) compared to castle (mound) type bowers and that hindbrain features evolve rapidly and independently of phylogeny among castle-building species. Using immediate early gene expression, we confirmed a functional role for hindbrain structures during bower building. Comparisons of bower building species in neighboring Lake Tanganyika show patterns of neural diversification parallel to those in Lake Malawi. Our results suggest that mosaic brain evolution via alterations to individual brain structures is more extensive and predictable than previously appreciated.

## Introduction

Animal behaviors vary widely, as do their neural phenotypes[1]. Evolutionary neuroscience identifies how the brain diversifies over time and space in response to selective pressures [2]. A key goal of evolutionary neuroscience has been to identify whether brain structures evolve independently (“mosaically”) or in tandem with each other as they reflect key life history traits, especially behavior [3-6]. While a number of studies have linked variation in brain structure with other traits across evolutionary time [2, 7-9], it remains unclear whether or not this variation is predictable. Specifically, when similar behavioral traits evolve among two or more species, do their neural bases evolve correspondingly? If parallel brain evolution is predictable then it may be possible to understand general principles of neural organization and function across animals. This would expand our ability to manipulate brain function, but if this is not true, new strategies will be needed to reveal the mechanisms of brain evolution.

Fishes, as both the most speciose (50% of extant vertebrates) and most varied vertebrate radiation [10] offer opportunities to answer these questions. Fish species live in diverse ecological, sensory, and social environments and have evolved elaborate variations in neural structure and function from a common basic ground plan [11] making rapid and variable diversification of brain structures a broad and general feature of their evolution [10].

The cichlid fishes of Lake Malawi, Africa offer a particularly striking model of these patterns of diversification. Although geologically young (less than 5 million years old), Lake Malawi contains at least 850 species of cichlids [12] that, based on molecular phylogenetic analyses, can be sorted into six diverse clades [13]: sand-dwelling shallow benthic species (287 species), deep benthic species (150 species), rock-dwelling ‘mbuna’ (328 species), the deep-water pelagic genus *Diplotaxodon* (19 species), the pelagic genus *Rhamphochromis* (14 species), and the sand-breeding pelagic ‘utaka’ species (55 species) [14]. These clades exhibit substantial variation in habitat use, visual sensitivity, diet, behavior, and coloration as well as craniofacial morphology and tooth shape, presumably arising through repeated divergence in macrohabitat, trophic specializations, diet, and coloration [15-17].

We studied the behavior, brain structures and bower construction of ~200 sand-dwelling shallow living benthic species with males that build species-specific mating nests, known as bowers, to court females in competition with conspecifics [17,18]. There are two basic bower types: 1) pits or depressions in the sand, and 2) castles, where sand is heaped into a volcano structure with showing species-specific differences in size and shape [17,18]. Pits and castles are constructed via differential scooping and spitting of sand. Pit and castle type bower building is innate and appears to be rapidly and relatively free from phylogenetic constraint; multiple sand-dwelling genera contain both pit and castle-building species [17]. Furthermore, phylogenomic analyses suggest evidence of the repeated evolution of bower types in Malawi cichlid phylogeny [17].

How is construction of different bower types reflected in brain anatomy? Do differences in brain structure evolve in tandem with bower building and, if so, how are they organized and functionally related to this behavior? Here we address these questions using neuroanatomical, phylogenetic, functional, and behavioral analyses and compare patterns of diversification in Lake Malawi to other East African cichlid radiations.

## Results

### Bower type predicts hindbrain volume

We compiled measurements of six brain regions (telencephalon, hypothalamus, cerebellum, optic tectum, olfactory bulb, and hindbrain) for 37 bower building species from a phenotypic data set comprising brain measurements for 189 cichlid species from East Africa and Madagascar [19]. For each brain region, we calculated a volume estimate using an ellipsoid model [20] and controlled for size differences between species by normalizing the results using mean standard length for each species (Supplementary material). Pairwise comparisons of normalized volumetric measures for each region revealed a significant difference in hindbrain volume between pit digging and castle building species (Kruskal-Wallis test; H = 13.87, 1 d.f., *p* = 0.00019) but not for any other brain structure (Table 1). Plotting structural volume against standard length reveals strong evidence for allometric scaling of the pit and castle hindbrain (Figure 1A). An analysis of variance (ANOVA) comparing the slopes of standard length compared to those of hindbrain volume reveals that pit and castle species significantly differ in this relationship (F = 25.04, 1 d.f., *p* = 1.82×10^-05^) as also confirmed by post hoc Tukey test (*p* <0.05).

**Table 1.** Allometric comparisons of brain structure volume. Results from comparisons of pit and castle species’ brain structure volumes (normalized by standard length) using Kruskal-Wallis test. ^***^ indicates *p* < 0.001.

| Structure | Chi-squared statistic (H) | *p*-value |
| --- | --- | --- |
| Hindbrain | 13.874 | 1.9x10<sup>-4***</sup> |
| Telencephalon | 0.060 | 0.807 |
| Cerebellum | 0.046 | 0.830 |
| Olfactory bulb | 0.054 | 0.817 |
| Hypothalamus | 0.001 | 0.975 |
| Optic tectum | 0.790 | 0.374 |

**Fig 1.**
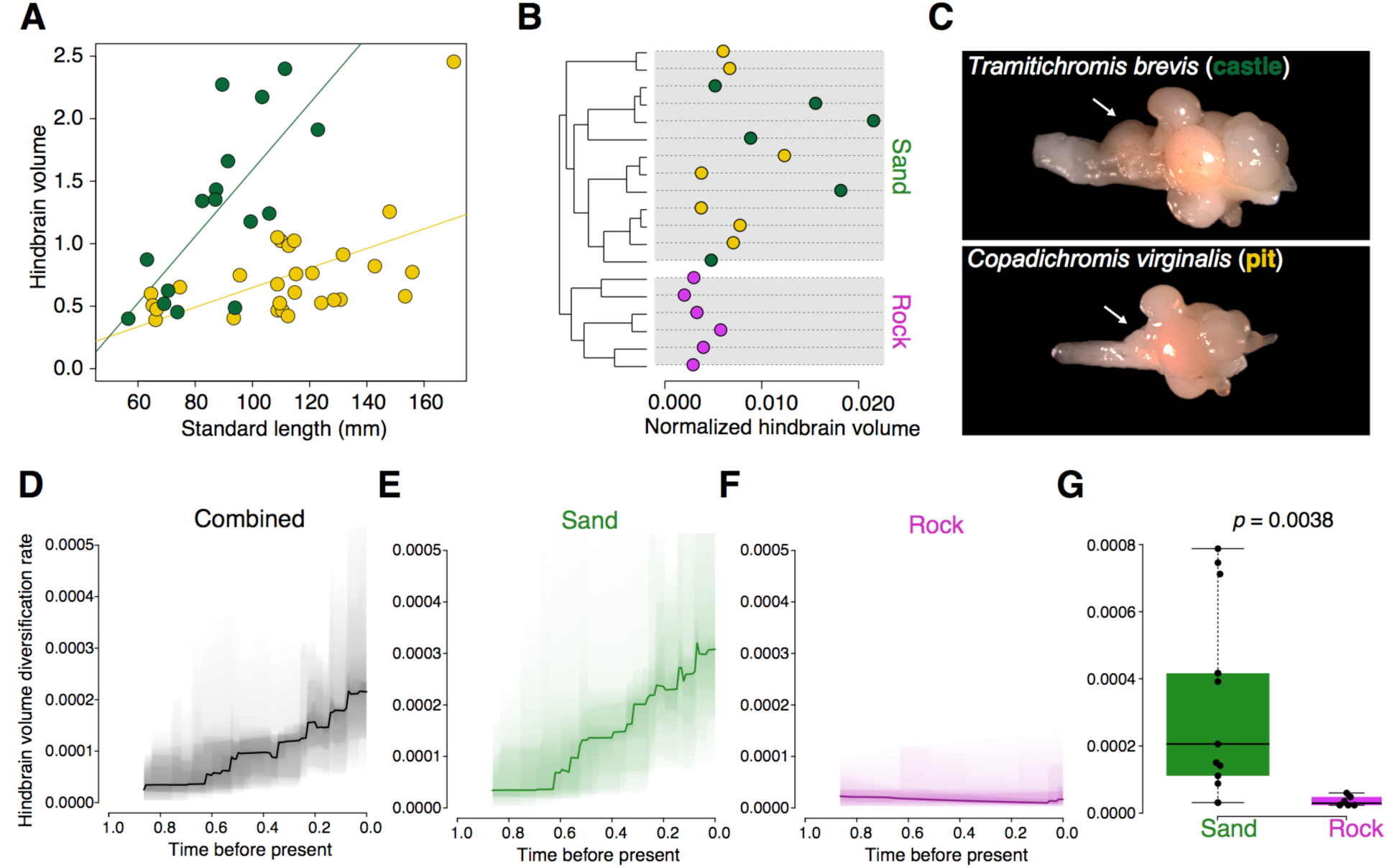
Evolution and diversification of hindbrain volume among Malawi cichlids. **(A)** Scatterplot comparing standard length and hindbrain volume for castle building species (green) and pit digging species (yellow). **(B)** Maximum likelihood phylogeny for 19 Lake Malawi sand and rock species. Hindbrain volumes normalized by standard length are represented by dots (castle = green, pit = yellow, rock = purple; for species labels and node support see Fig. S2). **(C)** Representative photos of whole brain and hindbrain (indicated by arrows) size for the castle species *Tramitichromis brevis* and the pit species *Copadichromis virginalis*. **(D-F)** Normalized hindbrain volume diversification rates for rock and sand **(D)**, sand alone **(E)**, and rock alone **(F)**.

### Hindbrain volume varies independent of phylogeny

If hindbrain volume evolved mosaically, as suggested by our allometric analyses, then pairwise tests of phenotypic variation controlling for phylogeny should reveal a lack of similarity among closely related species in the size of this brain region. Phylogenetic ANOVA using a genome-wide maximum likelihood phylogeny [21] revealed significant variation between pit and castle species independent of relatedness (F = 13.18, 1 d.f.,*p* < 0.05). Plotting the distribution of volumetric measures along the full tree demonstrates this pattern (Figure 1B). We next calculated more explicit measures of the phylogenetic signal using Pagel’s λ (Pagel 1999) and Blomberg’s *K* [22]. Pagel’s λ ranges from 0 to 1 with values closer to 1 representing stronger phylogenetic signal while values of Blomberg’s *K* < 1 suggest less phylogenetic signal in the trait. Both measures indicated a lack of phylogenetic signal among the species tested (λ = 0.25; *K* = 0.29), suggesting that hindbrain volume may be rapidly evolving but not correlated to phylogeny [22].

Since recently evolved clades such as Lake Malawi cichlids are prone to processes such as introgression and incomplete lineage sorting that make resolving phylogenetic relationships difficult [12], we re-performed the phylogenetic ANOVAs and tests of phylogenetic on non-overlapping genomic windows each containing 10,000 SNPs (1,029 windows; Figure S1A). We reasoned that if the patterns revealed by the genome-wide tests held in a substantial number of these windows then the observed results should be robust to alternative phylogenetic scenarios among the species sampled. Applying phylogenetic ANOVAs across the windows we found that all tested yielded a *p*-value less than 0.05 with the median of 0.006 (Figure S1B). Similarly, median values of phylogenetic signal were less than 1 across all windows (median λ = 0.011; median *K* = 0.728; Figures S1C-D). Of note, the distributions of *K* and λ are multimodal, suggesting that multiple phylogenetic patterns may still exist within the genomes of these species. This is reflective of the genomic complexity found among Malawi cichlids mentioned above and may have been captured by the relatively large window size used for these tests. Nonetheless, taken together these results support a model in which hindbrain volume, at least among the species tested, is associated with a deficit of phylogenetic signal.

### Castle building is associated with increased rates of hindbrain diversification

We analyzed rates of trait diversification between rock and sand Malawi cichlid species with a whole-genome phylogeny using Bayesian Analysis of Macroevolutionary Mixtures (`BAMM`) v.2 [23, showing that hindbrain volume has increasingly diversified since the split between rock and sand lineages around 800,000 years ago (Figure 1D). Separating the overall phylogenetic tree into rock and sand clades revealed extremely different rates of diversification between these two groups with the sand group (including bower builders) displaying a 7.4-fold faster rate of phenotypic diversification (Figure 1E-F).

Given this rapid evolution, we hypothesized that hindbrain diversification rates would be greatest in more recently diverging clades. Consistent with this hypothesis, the model with the best rate shift configuration in `BAMM` included two significant shifts in younger bower-building subclades within the sand-dwelling lineage (Posterior probability = 31.58; Figure S2). We further found that three of the four best shifts accounting for the majority of posterior probabilities sampled within the 95% credible shift set significant shifts on at least one of these branches (Figure S2). These subclades were enriched for castle building species, including one that was entirely composed of castle-builders, suggesting that the evolution of castle building is associated with increased diversification rates and trait values for hindbrain size. The phylogenetic analyses conducted here, and previously [17], suggest that the construction of pit bowers may be ancestral from which castle building may have arisen multiple times. We then hypothesized that hindbrain volumes among pit digging species should be more similar to those of the rock lineage while castle building species should differ significantly. Indeed, hindbrain volumes of pit digging and rock dwelling species are statistically indistinguishable (Kruskal-Wallis test; H = 0.66, 1 d.f., *p* = 0.42) while volumes of castle building and rock dwelling species differ substantially (Kruskal-Wallis test; H = 16.74, 1 d.f., *p* = 4.28 × 10^-5^). Taken together these results demonstrate that hindbrain volume is evolutionary labile and appears to have rapidly diversified in concert with bower type, suggesting extensive mosaic evolution of this structure within the sand-dwelling lineage.

### Variation in hindbrain architecture and connectivity among pit and castle species

To identify whether or not variation in hindbrain volume is also associated with cytoarchitecture and/or connectivity in that structure, we compared the neuroanatomy of two approximately size matched species - *Copadichromis virginalis* (CV; pit-digging) and *Mchenga conophoros* (MC; castle building, smaller hindbrain compared to TB) - and a castle builder with a comparatively elaborated hindbrain, *Tramitichromis brevis* (TB; castle building, larger hindbrain compared to MC). Diversification of the hindbrain in other fish species is associated with expansion of the vagal lobe in the dorsal medulla, a key center for taste sensation and processing that receives projections from sensory structures involved in gustation [24]. We found that CV, MC, and TB displayed three grades of anatomical complexity in the morphology of both these nerves (vagal nerve complex) and in associated vagal sensory structures of the brainstem (oropharynx). The simplest morphology (Figure 2A-B), in *Copadichromis virginalis* (CV), was similar to that of most teleost genera examined, with a smooth epithelium to the posterior oropharynx with no specialized secondary structures such as masticatory pads or palatal organ. Correspondingly, the vagus nerve was relatively small, with a single main root, and the brainstem vagal complex was relatively small compared to the other species. In contrast, the oral apparatus and vagal complex of *Mchenga conophoros* (MC) was more highly developed. A distinct palatal organ (PO) was associated with the gill arches located most caudally and the oral palatal surface morphology was more convoluted (Figure 2C-D). Concomitantly, the vagal nerve was more developed showing multiple roots at the point of entrance to the brainstem and the central vagal complex was larger than in *C. virginalis*. More complex still was the oropharynx of *Tramitichromis brevis*. The posterior palate housed a large palatal organ whose epithelium was thrown into rough protruding papillae (Figure S3A-S3C). The surface protrusions exhibited numerous taste buds at their apex as revealed by staining for calretinin. The vagal nerve complex was elaborated with distinct rootlets penetrating each of the branchial arches (Figure 3b). The medulla was dominated by a large vagal lobe protruding from the dorsomedial surface.

**Fig 2.**
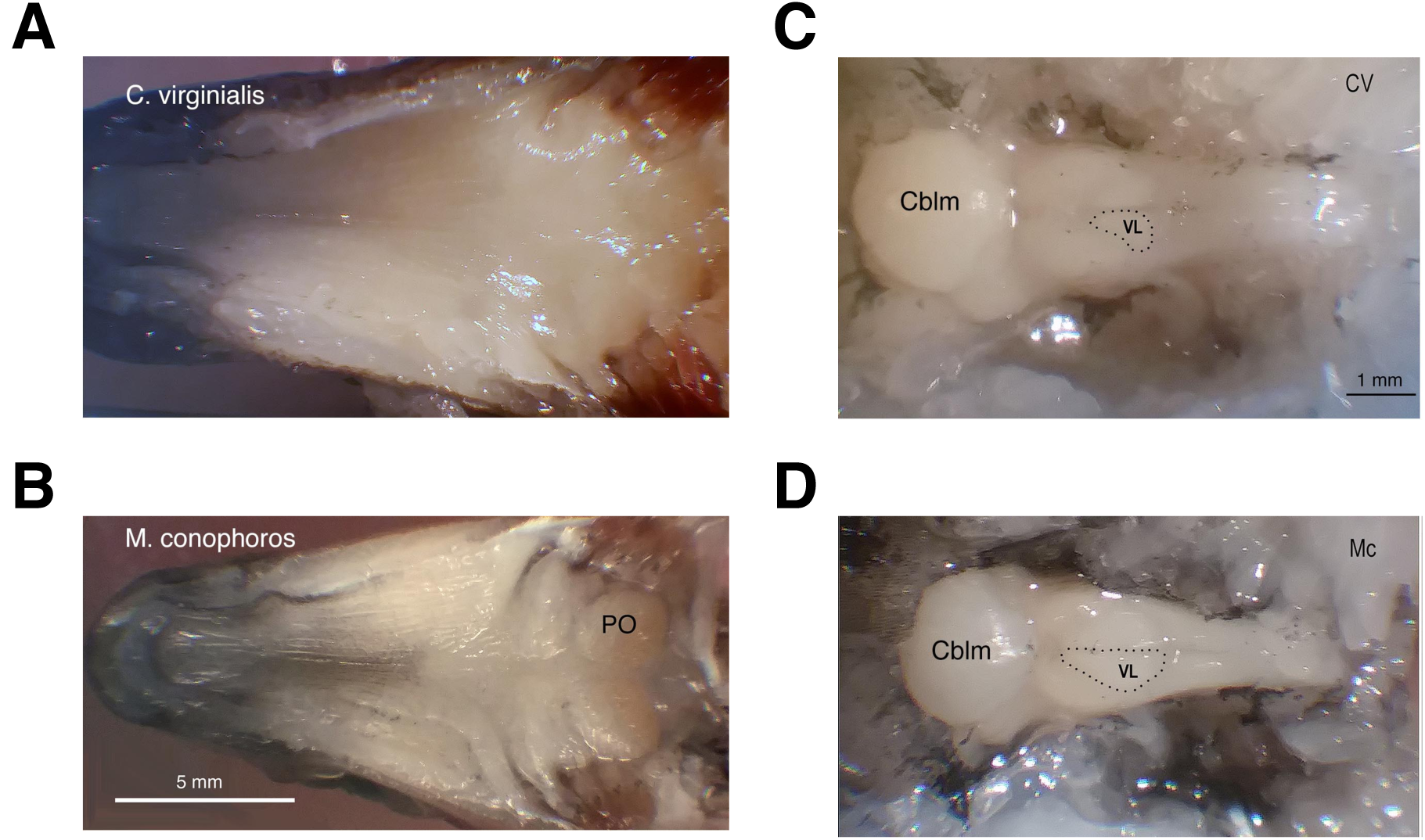
Oral cavity and brain morphology of *Mchenga conophoros* and *Copadichromis virginalis*. **(A)** The dorsal oral cavity of the pit-digger *Copadichromis virginalis* is comparatively simple and lacks a distinct palatal organ. **(B)** The dorsal oral cavity of the castle-builder *Mchenga conophoros* displays a more convoluted structure and caudal palatal organ labeled ‘PO’. **(C)** Photograph of *in situ* caudal brain structures of *C. virginalis*. The approximate location of the vagal lobe is outlined and labeled ‘VL’ and the cerebellum is labeled ‘Cblm’. **(D)** Photograph of *in situ* caudal brain structures of *M. conophoros*. Like in **(C)** the approximate location of the vagal lobe is outlined and labeled ‘VL’ and the cerebellum is labeled ‘Cblm’.

**Fig 3.**
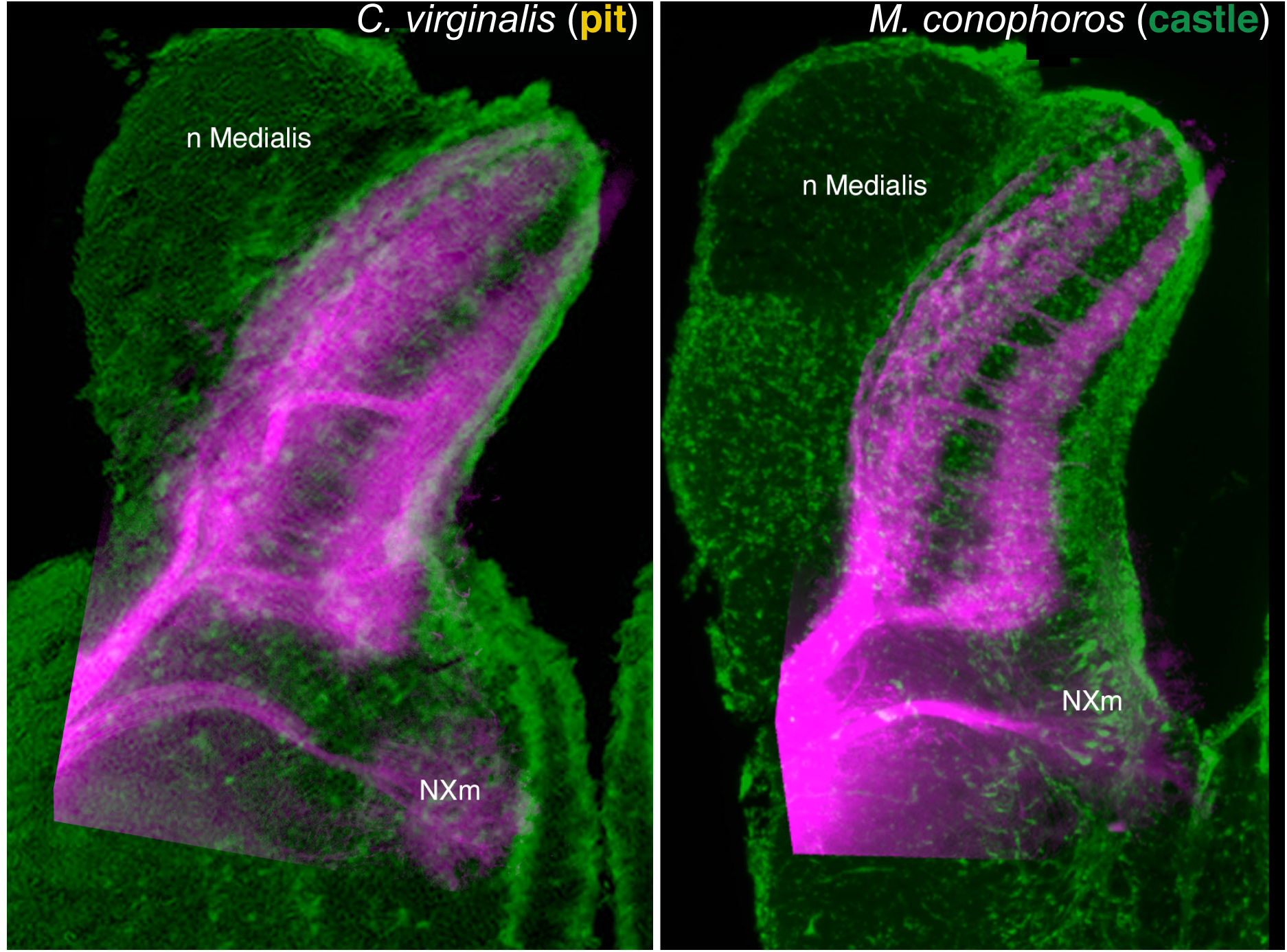
Vagus nerve termination. Composite micrographs comparing the laminar pattern of termination of vagus nerves in the vagal lobes of *C. virginalis* and *M. conophoros*. The sensory roots of the vagus nerve, labeled by DiI and rendered in magenta, terminate similarly in the two species: in superficial and deep layers leaving the central region void of terminals but crossed by fascicles running from the superficial root into the deeper layers. Green =Nissl staining inverted and rendered in green as if a fluorescent Nissl stain; magenta = diI label; image superimposed on the image of Nissl staining from another specimen.

Comparing sections through the vagal lobe complex of CV and MC showed differences in elaboration of the central vagal complex corresponding to the differences in oral anatomy (Figure 4, S3-4). In the caudal half of the medulla, the vagal lobe of CV showed a small, albeit clearly laminated vagal lobe. Numerous densely-packed small cells lay along the medial, ventricular face of the lobe at all levels. At the caudal end of the molecular layer of the nucleus medialis, two relatively superficial layers of small neurons were evident. Two or more relatively poorly defined layers of cells lay between these superficial layers and the periventricular layer. Although the overall size of the vagal lobe of MC was larger and more highly structured than that of CV, the essential laminated structure was similar. The vagal lobe of TB was larger still. In both TB and MC, a layer of densely packed small neurons lined the ventricle while densely packed and slightly clumped aggregates of small neurons dominated the superficial layers. As in CV, several layers of neurons with no apparent organization formed the central portion of the lobes.

**Fig 4.**
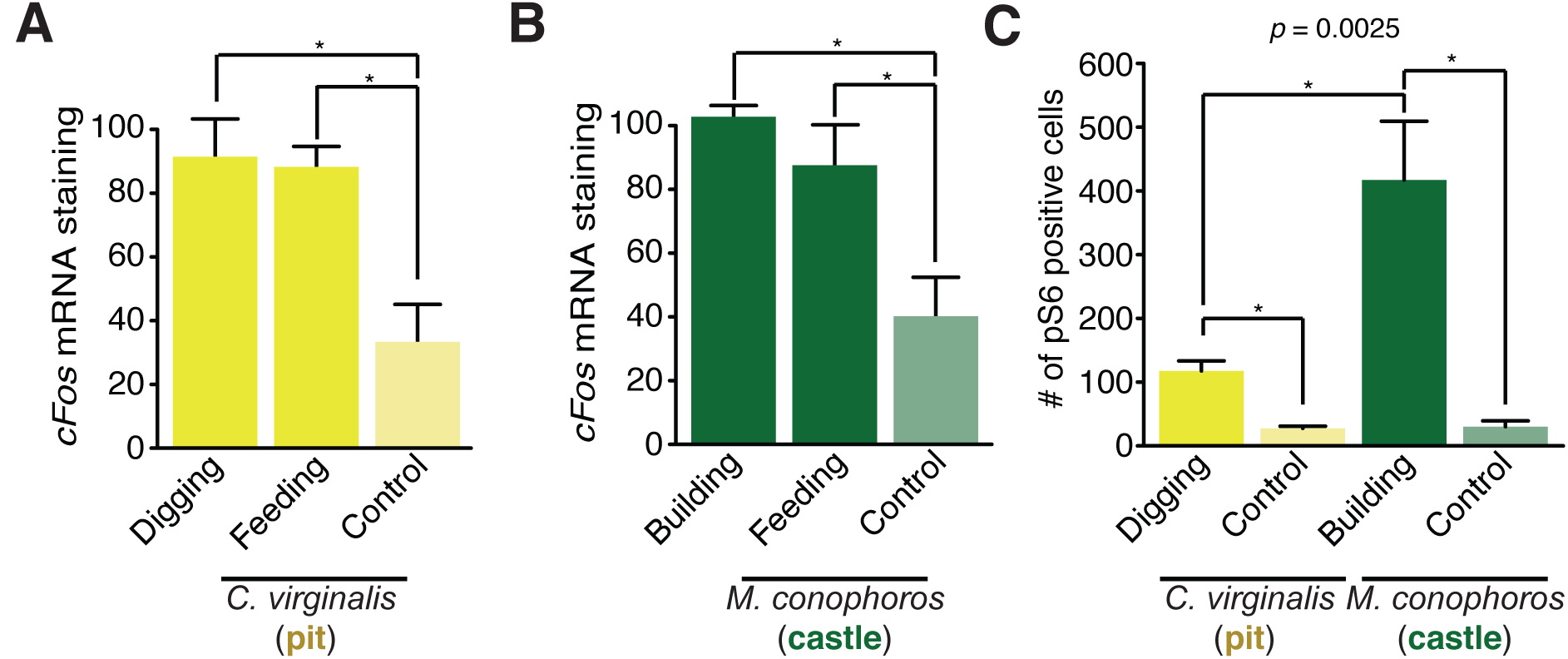
Molecular correlates of vagal lobe activity associated with bower building. **(A)** Barplot comparing *cFos* mRNA staining in MC across control, feeding, and building conditions across the VL. ^*^*P* < 0.05. **(B)** Barplot comparing *cFos* mRNA staining in CV across control, feeding, and building conditions across the VL. ^*^*P* < 0.05. **(C)** Barplot comparing pS6 positive cell abundance during bower building in *C. virginalis* and *M. conophoros*. ^*^*P* < 0.05.

The central projections and motor neurons of the vagus nerve were identified in all three species by post-mortem application of diI to the vagus nerve roots just distal to their entrance into the brainstem (see Methods). As seen in Figure S4, the pattern of termination of the vagus nerve in all three species appears similar. In all species, the sensory root of the nerve entered the lobe ventrolaterally and divided into a superficial and deep root terminating respectively in the superficial half of the lobe and in the deeper one-third of the lobe with an intervening zone of neuropil relatively free of vagal endings. This terminal-free zone was, however, penetrated by numerous fascicles of the nerve running from the more superficial root into the deeper terminal layers.

The motoneurons of the vagus nerve lay in a periventricular position ventral to the vagal lobe proper, as in most other species of fish (e.g. catfish) [24]. This differs from goldfish and common carp where vagal motor neurons lie along the innermost layer within the vagal lobe itself [24, 25].

The palatal apparatus and attendant vagal lobe are proportionately larger in TB than in the other 2 species examined. Nonetheless, the basic projections of the vagus nerve, the nature of the vagal lobe lamination and the position of the vagal motoneurons appear similar to those of the other species. In TB a distinct glossopharyngeal nerve root was identifiable (Figure S4N) which terminated in a glossopharyngeal region (Figure S4N-O) midway between the facial lobe (unlabeled by DiI positive fibers in Fig. S4M) and the more clearly laminated vagal lobe (Figure S4P-S). Although this glossopharyngeal region appeared continuous with both the facial and vagal lobes, it is clearly distinct from the latter in terms of cytological organization as in some cyprinid species [24, 25]. The vagal lobe has superficial and deep terminal zones similar to those in the other species described, whereas the glossopharyngeal zone does not. The terminal zone in the glossopharyngeal zone was continuous only with the dorsal terminal zone of the vagal lobe; no deep terminal zone was apparent. Separate glossopharyngeal and vagal motor neuron pools were evident below the ventrolateral edge of the ventricle, each pool being situated beneath the respective terminal field zone within the overlying lobes.

### The vagal lobe is functionally involved in bower building and feeding

We used *cFos* mRNA expression as a proxy for neural activity to assess which vagal lobe neurons are activated during bower building. Comparing the castle-building species *M. conophoros* (MC) and the pit-digging species *C. virginalis* (CV), we measured *cFos* expression in the vagal lobe during bower construction and compared it to control males who were housed without sand or female conspecifics (Figure S5A-F). Cells in the vagal lobe of bower building MC males showed greater *cFos* expression than control (2.601-fold difference, *p* = 0.019, bootstrap 1-way ANOVA, n/group > 3) (Figure 4A). The level of *cFos* expression in the vagal lobe of CV also increased significantly (2.517-fold difference, *p* = 0.034). Likewise, *cFos* expression in the vagal lobe increased during feeding in both MC (2.254-fold difference, *p* = 0.046) and CV (2.438-fold difference, *p* = 0.033) (Figure 4B).

To measure vagal lobe activation during bower building we used *in situ* hybridization of *cFos* and immunohistochemical labeling of phosphorylated ribosomal protein S6 (pS6). Labeling of pS6 indicates recent neural activity and displays punctate staining, allowing for granular quantification in individual neurons [26]. Staining of pS6 revealed that the number of VL neurons activated during bower building differed significantly between CV and MC. As with *cFos*, pS6 cell labeling was robust throughout the vagal lobe of both CV and MC after bower building (Figure S6A-H). Quantification of pS6 labeling in the vagal lobe showed significant differences during bower building in both species (CV: 4.12 fold difference, *p* = 0.015; MC: 13.84 fold difference, *p* = 0.022). Moreover, we found that MC bower building males displayed an almost four-fold greater abundance of pS6 positive neurons compared to CV (3.68 fold increase, *p* = 0.04) (Figure 4C).

### Hindbrain volume has evolved in parallel with bower building in Lake Tanganyika

To ask whether parallel diversification of hindbrain volumes may have occurred in non-Malawi cichlids, we measured variation in hindbrain volumes in the cichlid radiations of Lakes Victoria and Tanganyika. We found that the Lake Victoria species sampled had hindbrain volumes similar to those of the rock-dwelling cichlids of Lake Malawi (Figure 5A) whereas normalized hindbrain volumes of Lake Tanganyika species were distributed similar to those of Malawi species (Kruskal-Wallis test, H= 0.043, 1 d.f., *p* = 0.84), providing evidence of increased hindbrain diversification within Tanganyika cichlids. This observation was notable given that Lake Tanganyika has a number of reported bower building species, in contrast with Lake Victoria where few, if any, are known. This suggests that hindbrain diversification has occurred where cichlids have evolved this specific behavior (Malawi and Tanganyika), but not in lakes where bower-building does not occur (Lake Victoria). Accordingly, differences in the hindbrain volumes of Tanganyikan bower building species compared to non-bower builders were comparable to those found between the Malawi rock and sand lineages (Figure 5A).

**Fig 5.**
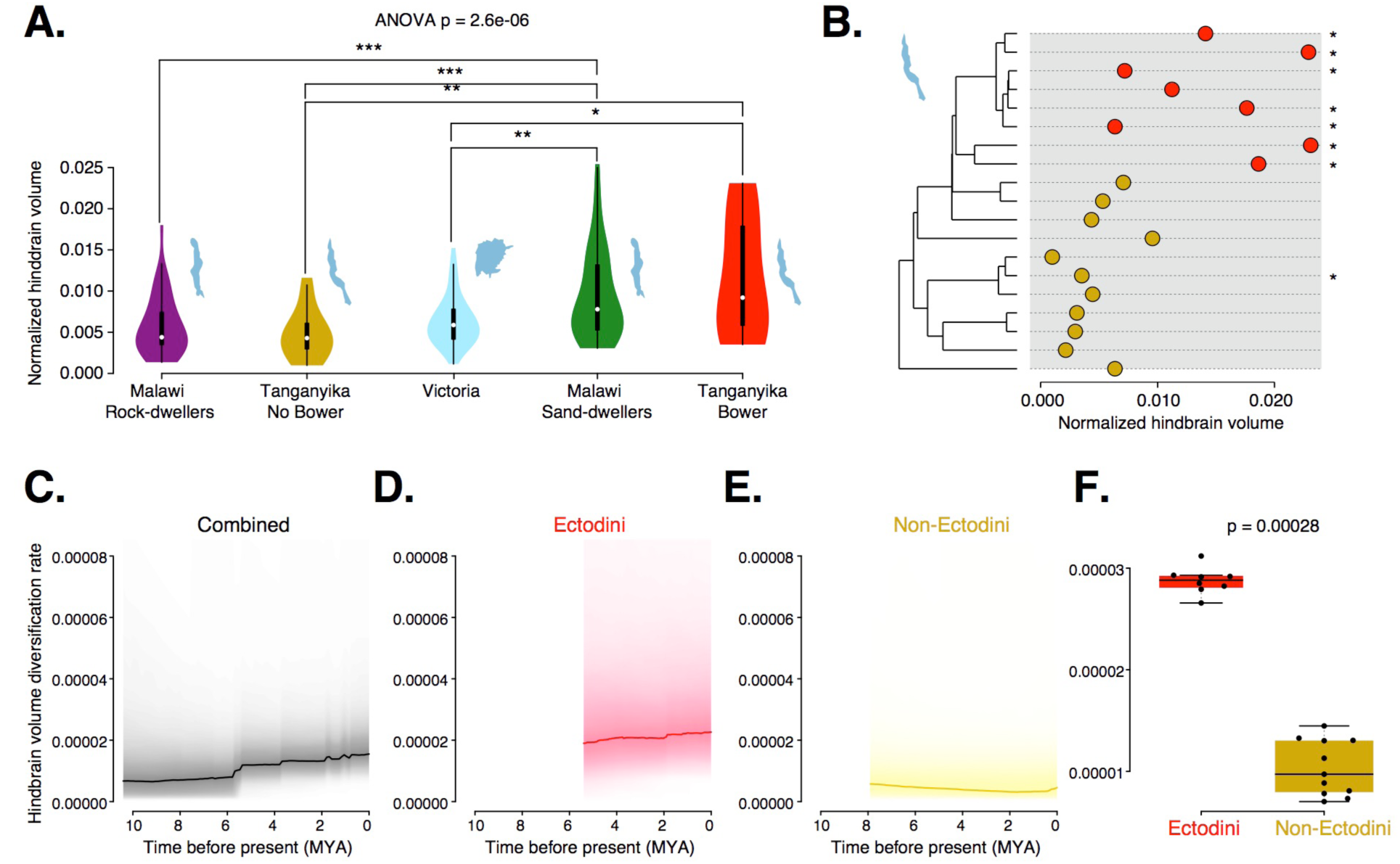
Evolution and diversification of hindbrain volume across East African Cichlids. **(A)** Violin plots of normalized hindbrain volumes for Malawi rock-dwelling species (n=47), Lake Tanganyika non-bower building species (n=14), Lake Victoria species (n=55), Malawi sand-dwelling species (n=37), and Tanganyika bower building species (n=12). Lakes are represented by cartoon outlines accompanying their respective violin plots. **(B)** Maximum likelihood phylogeny of Lake Tanganyika species sampled with accompanying normalized hindbrain volumes. Red represents the Ectodini tribe, yellow represents non-Ectodini, stars indicate known bower building species (for species and node labels see Fig. S10) **(C-E)** Normalized hindbrain volume diversification rates for Ectodini and non-Ectodini **(C)**, Ectodini **(D)**, and non-Ectodini **(E)**. **(F)** Boxplot comparing per-species normalized hindbrain volume diversification rates between Ectodini and non-Ectodini tribes.

Phylogenetic visualization revealed that hindbrain diversification within Lake Tanganyika was largely limited to the Ectodini tribe that has a number of bower building species (Figure 5B). We found that in this phylogenetic grouping of bower builders, in contrast to Lake Malawi cichlids, there was little hindbrain evolution independent of the Lake Tanganyika phylogeny (ANOVA (F = 9.79, 1 d.f., *p* = 0.12). Similarly, we identified significant amounts of phylogenetic signal in hindbrain volume (*K* = 0.53, *p* = 0.035; λ = 0.65, *p* = 0.048).

Analyses of diversification rates using `BAMM` showed hindbrain diversification has increased within the Lake Tanganyika Ectodini tribe as compared with non-Ectodini species (Figure 5C-E). Our data support a model of parallel diversification of hindbrain volume associated with bower building in both Lake Malawi and Lake Tanganyika, but with substantially different evolutionary histories associated with the trait in each lake.

## Discussion

We show here that hindbrain volume increases in tandem with the behavioral evolution of castle-type bower building, both within the dozens of species sampled from Lake Malawi here and also as compared to species in Lake Tanganyika but not Victoria, which seems to contain no known builders. Despite previous results identifying mosaic evolution of the brain, whether this may occur repeatedly given common ecological, evolutionary, or phylogenetic pressures is unclear. Our findings suggest that in certain scenarios brain evolution may proceed in a predictable manner, as previously proposed for systems such as bony fish [10] and vocal learning birds and mammals [8], but has rarely has this been tested explicitly in a controlled phylogenetic context. However, resolving phylogenetic relationships among Malawi cichlid species is difficult given their close genetic relationships and complex histories of gene flow and incomplete lineage sorting [14]. Therefore, future work attempting to disentangle further the differential roles of ecology, behavior, and evolution on brain morphology in Lake Malawi should include wider, more targeted sampling from the phylogeny in addition to comprehensive collection of demographic traits for each species. Sequencing more Malawi species would allow useful phylogenomic analyses of genetic association and identification of the roles of gene flow and/or incomplete lineage sorting in the context of brain evolution.

Our results also indicate that variation in hindbrain size is associated specifically with diversification of the vagal lobe, a key gustatory region of the fish hindbrain. The vagal lobe differs significantly between pit and castle species in size and structure, but not connectivity. This contrasts starkly with observations in cyprinid fish, another teleost clade with extensive vagal lobe diversification. Among typical food-sorting cyprinids that have a palatal organ, such as goldfish and carp, the vagal lobe is highly laminated with clear motor and sensory zones that extend dorsally in parallel along the deep and superficial layers of the lobe [24, 25, 27]. Similarly, the vagal lobe of *Heterotis niloticus*, an unrelated fish of the order *Osteoglossiformes*, has a laminated vagal lobe with sensory layers overlying motor layers [28]. In fish species that lack an elaborate palatal organ, such as catfish and zebrafish, the vagal lobe is cytologically simpler with less obvious lamination and a motor nucleus which is restricted to a sub-ventricular area with little penetration into the lobe itself [29]. Malawi cichlid hindbrain diversification is an intermediate between the vagal lobes of food sorting and non-food sorting cyprinids. While the vagal motor nuclei of pit and castle species is similar in location to that of catfish, histology and DiI labeling indicate that castle building species tend to show increases in vagal lobe size, lamination, and manner of termination of primary vagal sensory fibers. These patterns of vagal lobe elaboration are supported by the presence of palatal organs in just the castle building species sampled. Furthermore, our studies of immediate early gene expression and ps6 abundance during behavior reveal not only that the vagal lobe is involved in both bower building and feeding, but that there are detectable species differences in neuronal activity during bower building. Such differences may merely reflect differences in the degree to which oral manipulation of substrate is necessary for the different types of bower: hole versus structure.

Evidently, the largest differences in hindbrain structure and function among bower builders are determined mostly by activity (via behavioral state) and size, but only moderately by changes in connectivity and histological structure. Any functional differences in the vagal lobe associated with pit and castle bower types, then, likely arise through both variations in developmental patterning and, in adulthood, modulation of vagal lobe function. It does not appear that bower building is associated with the evolution of novel, behavior-specific hindbrain circuits. Support for this comes from previous work showing that ecologically-relevant differences in forebrain size among rock- and sand-dwelling Malawi cichlids arise from a common blueprint that is differentially modified by patterning genes [30]. Similarly, the hindbrains of the sand-dwelling cichlids analyzed here appear to have a relatively conserved brain bauplan at base, but with substantial elaboration in size and modulation among species. The high degree of relatedness and recent evolution of Malawi cichlid species likely constrains the phenotypic possibilities available for the evolution of brain and behavior. The significant increase in diversification rate of the hindbrain among castle-builder clades in both Malawi and Tanganyika suggests that behavioral evolution in these lakes is likely supported by neural variation producing bigger brain structures that generate more activity.

The degree of phenotypic predictability displayed by the hindbrain in Malawi cichlids suggests that, given a common phylogenomic context, diversifying species may present some degree of commonality in the evolution of neural and behavioral traits in response to similar pressures. Of particular interest will be the study of how evolution acts on conserved and novel genes and genetic networks to regulate the brain and behavior across evolutionary distances, potentially revealing common principles of the evolution of behavior.

## Methods

Fish were bred, housed, and maintained at Stanford University following established Stanford University IACUC protocols.

Volumetric analyses were performed using measurements from a phenotypic data set comprising brain measurements for 189 cichlid species from East Africa and Madagascar [19]. Phylogenetic comparisons were conducted in R [38]. The packages SNPhylo [31] and ape [32] were used to construct ultrametric phylogenies for the Lake Malawi species from whole genome SNP data [21]. A previously published phylogeny was used to infer phylogenetic relationships of the Lake Victoria and Tanganyika species [35]. Phylogenetic ANOVAs were performed with the package geiger [38] while phylogenetic signal and trait diversification rates were inferred from the packages phyloSignal [40] and BAMMtools [23], respectively.

Oropharyngeal anatomy was assayed by scanning electron microscopy of the oral cavities of *C. virginalis* (CV), *M. conophoros* (MC), and *T. brevis* (TB) using a Hitachi S-3400N VP scanning electron microscope. Taste buds and innervation patterns were identified by immunocytochemical labeling using antisera against human calretinin (CR 7697; AB_2619710) and acetylated tubulin (Sigma T7451; AB_609894).

DiI tracing was performed on CV, MC, and TB by placing DiI crystals on either the vagus nerve root or the surface of the vagal lobe. After a diffusion period of 1-6 months the brains were sections at 75-100 um on a vibratome and then imaged.

Behaviorally-induced neural activity was assessed by running CV and MC males through one of three behavioral paradigms: bower building, feeding, and control. For ISH we used RT-PCR to amplify a portion of coding sequence from *cFos* (NM_001286320), and subcloned products into pCR-TOPO4 (Life Technologies). *cFos* forward, 5’-AAT TGG ATC CAA GCC CAG ATC TTC AGT GG-3’; *cFos* reverse, 5’-AAT TGA ATT CAT AGC CCT GTG ATC GGC AC-3’. Abundance of pS6 was inferred by immunohistochemical labelling with with rabbit anti Phospho-S6 Ser244 Ser247 (ThermoFIsher Scientific 44-923G, RRID AB_2533798). ISH and immunohistochemical labeling were imaged in FIJI and quantified with custom scripts in R.

Additional information can be found in the supplemental experimental methods.

### Vertebrate animal use

All animal experiments were approved by the Stanford University Animal Care and Use Committee (protocol number APLAC-28757).

## Data availability

All sequencing data in support of the findings of this study have been deposited in the Short Read Archive under accessions SRR6314224, SRR6314225, SRR6314226, SRR6314228, SRR6314230, SRR6322515. All morphometric, phylogenetic, and histological data are available as electronic supplements to this manuscript.

## Declaration of interests

The authors declare no competing interests.

## Acknowledgements

We thank Beau Alward, Austin Hilliard, Patrick McGrath, and Hunter Fraser for helpful comments on earlier drafts of the manuscript. We thank Robert Huber for sharing raw brain measurement data and Catherine Wagner for the Tanganyika and Victoria species phylogeny. This research was supported by awards from the NIH to RDF and JTS (R01GM101095), RDF (NS034950), from the Stanford Bio-X Bowes Fellowship (RAY) and a Stanford CEHG Fellowship (RAY). The authors have no competing interests.

## Contributions

RY conceived of the study, designed the study, carried out behavioral and molecular experiments, performed all phylogenetic and statistical analyses, and wrote the manuscript. AB performed behavioral assays and *in situ* hybridization and immunohistochemical staining protocols. KA, CP, and JTS collected and processed tissue samples for next generation sequencing and identified genetic variants for phylogeny construction. TF performed the neuroanatomical comparisons of Malawi cichlid hindbrain structure and helped draft the manuscript. RDF provided funding, aided in design and coordination of the study and helped write the manuscript.

